# Evolution of plasticity and character displacement in a fluctuating environment

**DOI:** 10.1101/2024.12.02.626345

**Authors:** Luis-Miguel Chevin, Lakshya Chauhan

## Abstract

Species that compete for the same resource may undergo ecological character displacement (CD), where the phenotype of each species evolves to deviate from its optimum in the absence of competition. As natural habitats are rarely static, interspecific competition is likely to occur in environments that fluctuate over time. Such environmental fluctuations can in turn cause the evolution of phenotypic plasticity for traits mediating the competition. However, the interplay of environmental fluctuations, evolution of plasticity, and character displacement, has been little investigated. We use a quantitative genetic model to study theoretically how a randomly fluctuating environment and evolution of plasticity influence the outcome of ecological CD between two competing species. We show that environmental fluctuations make the conditions for CD more stringent, requiring stronger competitive selection relative to stabilizing selection. This occurs because environmental fluctuations reduces the average population size, and thereby competition intensity. Evolving plasticity can restore CD by buffering the impact of environmental fluctuations through phenotypic tracking, to a degree that depends on environmental predictability. Somewhat paradoxically, competition that favors phenotypic divergence among species can cause convergence in plasticity, when this reduces the load caused by fluctuations in phenotypic divergence. Our results shed light on how competition and plasticity influences evolution of the fundamental niche in a fluctuating environment.

## Introduction

Species in their natural habitats need to adapt both to the abiotic challenges imposed by their physical environment, and to interacting species in their community. In the ecological literature, these biotic and abiotic components of the environment relate to the classical concepts of the fundamental versus realized niche (Hutchinson 1957; Holt 2009). The range of abiotic environments over which a species can sustain population growth, when alone and at low density, and thus persist without inputs from a more productive source population, defines the fundamental niche (Hutchinson 1957; Pulliam 1988; Holt 2009). This range can be reduced by antagonistic interactions such as competition, leading to a smaller realized niche (Hutchinson 1957; Holt 2009). From an evolutionary standpoint, the biotic and abiotic environment act as different, sometimes conflicting, sources of selection, the relative importance of which may change along environmental gradients. Not only does the number of interacting species change along abiotic gradients, e.g. across latitude and altitude (von Humboldt and Bonpland 1807) or salinity (Williams 1998), but the intensity and even the type of interactions may also change (Maestre *et al*. 2009). An important consequence of such variation in the strength of ecological interactions along environmental gradients is that these interactions may influence how species adapt to the abiotic environment, thus causing evolution of the fundamental niche.

One of the best-studied examples of evolution caused by interspecific interactions is ecological character displacement (hereafter character displacement, or CD) resulting from competition between species (Brown and Wilson 1956; Grant 1972; Schluter 2000). When sympatric species compete for the same resource, then any trait involved in their interaction can evolve away from its optimum in the absence of a competitor, thereby minimizing the detrimental effect of competition on fitness. This process has been documented empirically in a number of systems, mostly in vertebrate animals (Pfennig and Murphy 2000; Grant and Grant 2006; Kirschel, Blumstein and Smith 2009; Stuart *et al*. 2014), and to a lesser extent in plants (Kooyers, James and Blackman 2017). Evidence for CD also comes from the reciprocal phenomenon of character release, whereby traits or niches evolve after interspecific competition has been relaxed (Grant 1972; Robinson and Wilson 1994; Dayan and Simberloff 1998; Bolnick *et al*. 2010).

The influence of competition on evolution is even more pervasive when species need to adapt to a changing environment. In fact, some of the most emblematic examples of CD occur in environments that fluctuate substantially over time, leading to variation in the direction and strength of selection. For instance in Darwin’s finches on the Galapagos island of Daphne Major, invasion of a competitor species (*Geospiza magnirostris*) more efficient at cracking hard seeds caused the evolution of ecological character displacement towards reduced beak size in the local species *Geospiza fortis* (Grant and Grant 2006). Interestingly, this trait was also shown to be under strongly fluctuating selection in this population, where variation in the precipitation regime leads to drastic changes in the relative abundances of different seed types (Grant and Grant 2002). Such fluctuating environments, when partly predictable, can in turn cause the evolution of phenotypic plasticity, whereby a single genotype can express different phenotypes depending on the environment (Gavrilets and Scheiner 1993a; Tufto 2000; Lande 2009). In fact, phenotypic plasticity has been invoked as a powerful mechanism for character displacement (Pfennig and Pfennig 2009), as demonstrated for the dietary polyphenism of spadefoot toad tadpoles (Pfennig and Murphy 2000). While the most direct mechanism for plastic CD would involve phenotypic responses to competitor abundance, plastic responses to resources themselves, or to an abiotic environmental variable that accurately predicts selection on the trait mediating competition, should also influence the outcome of CD in a fluctuating environment. When this occurs, and if evolution of plasticity in response to the abiotic environment can be altered by the presence of competitors, this will have cascading consequences on environmental tolerance breadth (Chevin, Lande and Mace 2010; Lande 2014), such that competition will cause evolution of the fundamental niche (Holt 2009). However this question has received surprisingly little attention from evolutionary theory.

Early theory on CD in a constant environment highlighted the importance of the relative strengths of disruptive selection caused by competition, versus stabilizing selection resulting from intrinsic adaptation to the environment (Slatkin 1980; Taper and Chase 1985; Taper and Case 1992; Doebeli 1996a). More recent work has allowed the environment to vary over space, to investigate how interspecific competition influences local adaptation and geographic ranges of species that are spread over environmental gradients (Case and Taper 2000; Goldberg and Lande 2006). One of the key findings from this work was that eco-evolutionary dynamics play a critical role in this process. Indeed as a species becomes more maladapted, its abundance is likely to decrease, leading it to exert weaker competition, and thus cause weaker selection, on the other species. Such eco-evolutionary dynamics and feedbacks are also likely to play out in a temporally (rather than spatially) variable environment.

Phenotypic plasticity has been included in a few models of character displacement. Goldberg and Price (2022) recently extended their earlier model (Goldberg and Lande 2006) to allow for phenotypic plasticity, but without letting plasticity evolve, and without environmental fluctuations. The interplay of phenotypic plasticity and competition was also addressed in the more specific context of predation risk by Peacor *et al*. (2006), but without an explicit framework for phenotypic evolution (such as quantitative genetics), and assuming a temporally constant environment. In addition, this earlier theory mostly relied on numerical simulations, precluding the generality that can be gained from an analytical treatment of eco-evolutionary processes. Here, we instead focus on a temporally fluctuating environment (as done previously by Johansson 2008, but without plasticity), and use a combination of simulations and analytical results to yield deeper eco-evolutionary insights. We ask: (i) How the conditions for CD in a constant environment depend on the interplay between demography and evolution; (ii) How environmental fluctuations modify these conditions for CD; (iii) How evolution of phenotypic plasticity alters the outcome for CD, and the role of environmental predictability in this process; and (iv) Whether CD can exist for plasticity itself (e.g. reaction norm slope), beyond its effect on expressed phenotypes. We explain how a fluctuating environment and evolving phenotypic plasticity modify the conditions for CD, and demonstrate that competition can cause *convergent*, rather than divergent CD for its plasticity.

## Model

### Fitness and selection

We model two sympatric competing species that adapt to a variable environment via the same phenotypic trait. These species are denoted as 1 and 2, but most equations below are written in generic form (with indices *i* and *j*) that applies to any of these species. We assume discrete non-overlapping generations, as this makes plastic, evolutionary, and demographic responses to the fluctuating environment simpler to analyze. Under weak selection and population growth, discrete-generation models approximates continuous-time models well.

We follow Case and Taper (2000) and Goldberg and Lande (2006) in assuming that a single quantitative trait *z* influences both the density-independent component of population growth (intrinsic rate of increase), and the density-dependent component resulting from competition, within and between species. More specifically, the per-generation change in population size of species *i* (on the log scale) is

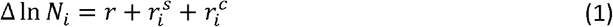

where the density-independent term

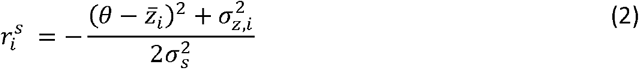

is largest when the mean phenotype 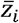 in species *i* matches the environmentally determined phenotypic optimum *θ*. This leads to stabilizing selection on 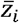, which is stronger when the width of the fitness peak is narrower (smaller 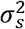). In line with previous models (Case and Taper 2000; Goldberg and Lande 2006), we assume that the phenotypic optimum *θ* is the same for both species, such that they would evolve to the same mean phenotype without competition. The competition component of population growth stems from logistic growth,

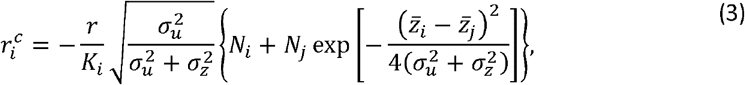

with *K*_*i*_ the carrying capacity of a hypothetical population with optimal phenotype in species *i*. Equation (3) shows that the population growth rate decreases with increasing population density *N*_*i*_ of the focal species, but also with the density *N*_*j*_ of the competing species. However, the latter is modulated by an Gaussian function of the mean phenotypic divergence between species 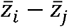, such that interspecific competition is stronger when species are more phenotypically similar. This captures Darwin’s (1859) classic hypothesis that “competition will generally be most severe between those forms which are most nearly related to each other in habits, constitution, and structure”, as verified experimentally with respect to phylogenetic similarity (Violle *et al*. 2011). The parameter 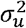 is the (squared) width of the resource utilization function, which determines the extent to which individuals overlap in their resource use, and thus how much they compete (Roughgarden 1979; Taper and Chase 1985). The term in the square root in eq. (3) arises from integration of the competition function among all interacting individuals within (and between) species (Case and Taper 2000; Goldberg and Lande 2006). It shows that the strength of density dependence is overall reduced when the phenotype distribution (with variance 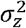) is broader than resource utilization, such that individuals overlap little in their resource use.

In line with standard quantitative genetics, we assume that the trait is polygenic, leading to a normal distribution of breeding values (heritable component) and phenotypes. Then using Lande’s equation (Lande 1976, 1982), the response to selection by the mean phenotype in species *i* is

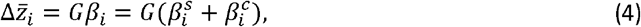

where *G* is the additive genetic variance of *z*, and the directional selection gradient *β*_*I*_ into a stabilizing selection component is decomposed

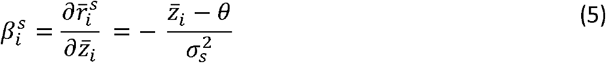

and a competition component

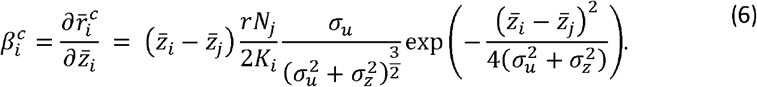

The first component of selection 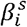 pulls the mean phenotype 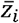 of the focal species towards the optimum determined by the environment, at a rate that depends on the squared width 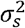 of the fitness peak. The second component 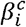 pushes 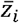 away from the mean phenotype 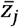 of its competitor species, at a rate that depends notably on the strength of density dependence, measured by 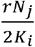. Taken in isolation, 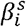 would lead to a stable equilibrium for 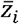 at the optimum θ (where 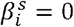 and 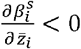, hence the term stabilizing selection), while 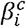 would lead to an unstable equilibrium for 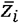 at the mean phenotype 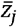 of the competing species (where 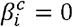 and 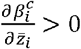 disruptive selection).

### Fluctuating environment and plasticity

To model the influence of a fluctuating environment on character displacement, we assume that an ε (e.g. temperature) causes linear changes in the optimum,

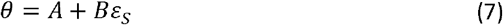

where *B* is the environmental sensitivity of selection (Lande 2009; Chevin, Lande and Mace 2010), and ε_*S*_ is the environment at the time of selection. The expressed phenotype may also depend on the environment through phenotypic plasticity. We assume developmental plasticity, such the expressed phenotype depends the environment ε_*D*_ experienced when the individual matured and the plastic response was fixed (Lande 2009). We also assume a linear reaction norm,

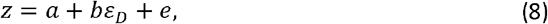

where the reaction norm intercept *a* and slope *b* are assumed to be normally distributed quantitative traits, with additive genetic variances *G*_*a*_ and *G*_*b*_, respectively, and covariance *G*_*ab*_.

Reaction norm slope *b* quantifies plasticity, while its intercept *a* is defined (without loss of generality) in a reference environment where it is uncorrelated with *b* (*G*_*ab*_ =0), which always exists with linear reaction norms (Gavrilets and Scheiner 1993b; Lande 2009). The normally distributed residual component of phenotypic variation *e* has mean 0 and variance *V*_*e*_ Under these assumptions, the selection gradients on reaction norm intercept and slope are

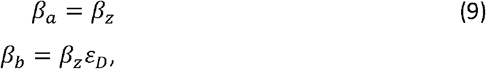

where *β*_*z*_ is the selection gradient on the expressed trait (difined *β*_*I*_ as for species *i* above) (Gavrilets and Scheiner 1993a; Lande 2009). Because the reaction norm parameters are not genetically correlated, their selection responses can be treated as univariate,

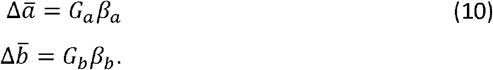

We assume that the environmental variable affecting both plasticity and selection fluctuates randomly over time, with a stationary normal distribution with mean 0 and variance 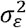, and positive autocorrelation that declines geometrically with time difference. In other words, ε_*t*_ undergoes a first-order autoregressive process (AR1), or an Ornstein-Uhlenbeck process sampled at discrete intervals(Karlin and Taylor 1981). We define *ρ* as the autocorrelation of ε_*t*_ between development and selection, separated by a time interval *τ* (lower than the generation time), and autocorrelation over any other time interval Δ *t* is 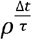.. If the competing species have different developmental times, then *ρ* is defined for species 1 (with delay *τ*_1_ between development and selection). Note that such stationary stochastic fluctuations differ from the non-stationary, random walk modeled by Johansson (2008) under a similar form of competitive selection.

### Mathematical analysis

To make the model analytically tractable, we relied on several assumptions, some of which we relaxed in simulations below. First, we mostly focused on the symmetric case where all demographic and selective parameters are identical in both species (i.e. same carrying capacity *K*, phenotypic variance 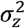, etc). This not only simplifies the system by halving its number of parameters and dynamical variables, but also corresponds to the most favorable case for character displacement to exist in both species, rather than competitive exclusion and maintenance of a single species with optimum mean phenotype. In addition, since the full eco-evolutionary system with joint demographic (eq. 1) and evolutionary (eq. 4) dynamics proved challenging to solve (or led to unwieldy results), our strategy was to derive the equilibrium population density assuming that the mean phenotype was at the optimum in both species 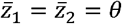, i.e. no CD) in eq. (1), and then used this equilibrium population density to search for evolutionary equilibrium using eq. (4), and thus identify threshold conditions for character displacement (CD).

### Simulations

The behavior of the model under scenarios of interest was explored by thorough simulations, using a program coded in Python3. These eco-evolutionary simulations consisted of joint recursions for demography (eqs. 1-3) and evolution (eqs 4-6), assuming constant genetic variance for simplicity (as in similar models by Case and Taper 2000; Goldberg and Lande 2006; Goldberg and Price 2022). This assumption amounts to considering that genetic variance has reached an equilibrium between mutation and stabilizing selection, and neglecting the additional noise caused by fluctuations of genetic variance around this equilibrium. The simulations took as input the initial values for the reaction norm parameters 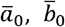 and population size *N*_0_, for each species. The scenarios we investigated differed regarding whether or not they included fluctuating selection (*B* = 0 vs *B* ≠ 0), initial plasticity 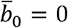 vs 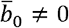), and evolution of plasticity (*G*_*b*_ = 0 vs *G*_*b*_ ≠ 0). Other parameters that varied (summarized in Table 1) were the variance and autocorrelation of environmental fluctuations (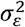 and *ρ*), and the delay between maturation and selection, which could differ between species (*τ*_1_ and *τ*_2_).

**Table 1:** Summary of variables and parameters of the model. All parameters are assumed to have the same values for both species (symmetric system), unless specified otherwise (notably for *ε*_*D*_ and *τ* in Figure 3).

| Symbol | Description | Value |
| --- | --- | --- |
| $N$ | Population size | - |
| $r$ | Maximum intrinsic growth rate | 0.1 |
| $K$ | Carrying capacity | 60000 |
| $\bar{a}$ | Reaction norm intercept | - |
| $\bar{b}$ | Reaction norm slope, quantifying plasticity | - |
| $\bar{z}$ | Expressed trait, accounting for plasticity | - |
| $\sigma_z^2$ | Phenotypic variance of expressed trait | - |
| $\sigma_e^2$ | Residual phenotypic variance for a given genotype in a given environment | 0.5 |
| $G_a, G_b$ | Additive genetic variances of reaction norm intercept and slope, respectively | 0.5, 0.045 |
| $G_{ab}$ | Additive genetic covariance between reaction norm intercept and slope | 0 |
| $\varepsilon_D / \varepsilon_S$ | Environment at development / selection | - |
| $\sigma_\varepsilon^2$ | Temporal variance of environment $\varepsilon$ | - |
| $\rho$ | Environmental autocorrelation between development ($\varepsilon_D$) and selection ($\varepsilon_S$) for focal species (species 1) | 0.5 |
| $\tau$ | Fraction of generation time between development and selection | 0.5 |
| $\sigma_s^2$ | Width ("variance") of stabilizing selection curve | 550 |
| $\sigma_u^2$ | Width ("variance") of resource utilization curve | 10 |
| $A, B$ | Intercept and slope of changes in the optimum phenotype with the environment | 5, 3 |

For each scenario, we conducted extensive parameter exploration with respect to the peak width for stabilizing selection 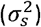 and the width of utilization function 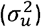.We used the package pypet (Meyer and Obermayer 2016), which allows efficient multiprocessing and facilitates parameter manipulation for multiple runs of a model. For each parameter combination in this exploration, the populations were left to evolve for >200,000 generations, and the last 10,000 generations were averaged over time to obtain the stationary values for the mean reaction norms elevation 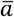 and slope 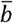 (plasticity), expressed trait 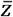, and population size *N*, for each species. The complete commented code along with execution instructions is hosted along with results at *ANONYMIZED*.

## Results

We wish to understand how a fluctuating environment and evolving phenotypic plasticity affect evolution of character displacement (CD) in two sympatric competing species. As a point of reference to characterize these effects, we start by analyzing a model with a constant environment and no plasticity. Our main argument focuses on symmetric conditions where both species are equivalent and interchangeable, such that most indexes for species can be dropped.

### CD depends on the relative strengths of stabilizing vs competitive selection

In a constant environment, an evolutionary equilibrium occurs when stabilizing selection exactly compensates for competition-mediated disruptive selection, such that 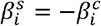 in both species (from equations 4-6). A trivial solution is where both gradients vanish, 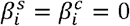 (for *i* = 1,2), such that the mean phenotype is at the optimum for both species and there is no 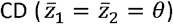. Another non-trivial solution with CD is straightforward to find in the symmetric case where phenotypic maladaptation and equilibrium population size are identical in both species. Then denoting as 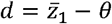 the deviation from optimum in species 1 (and thus 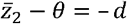 in species 2), solving for 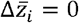 in eq. (4) assuming *d* ≠ 0 leads to

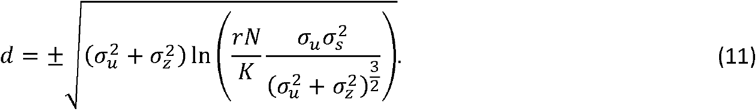

This solution is real when the term inside the logarithm is larger than 1, such that *d* is the square root of a positive number. This leads to a criterion for CD in terms of the width *σ*_*s*_ of the fitness peak,

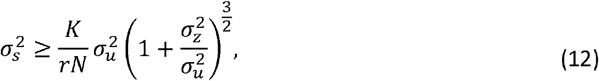

Equation (12) shows that CD can only occur when stabilizing selection is sufficiently weak relative to competitive selection (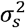 sufficiently large relative to 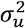). Stronger competitive selection, caused by either a larger (scaled) population density *N/K* or a narrower utilization curve (smaller 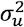, as long as 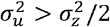) allows CD to exist in the face of stronger stabilizing selection. Nevertheless, the criterion in eq. (12) is a necessary but not sufficient condition for CD to exist. If both species start exactly at the optimum phenotype, then they remain “trapped” in the trivial equilibrium without CD, even for parameter values that could lead to CD. This trivial equilibrium is however unstable, and any minor deviation from it (caused by e.g. random genetic drift or a fluctuating environment, see below) eventually leads to CD when criterion (12) is met. This is illustrated in Figure 1A, where both competing species start with mean phenotypes close to the optimum *θ*, but slight deviations above and below this unstable equilibrium lead to phenotypic divergence and character displacement. Note also that the threshold in eq. (12) correspond to conditions at which *d* = 0 such that the equilibrium with CD disappears (bifurcation).

**Figure 1:**
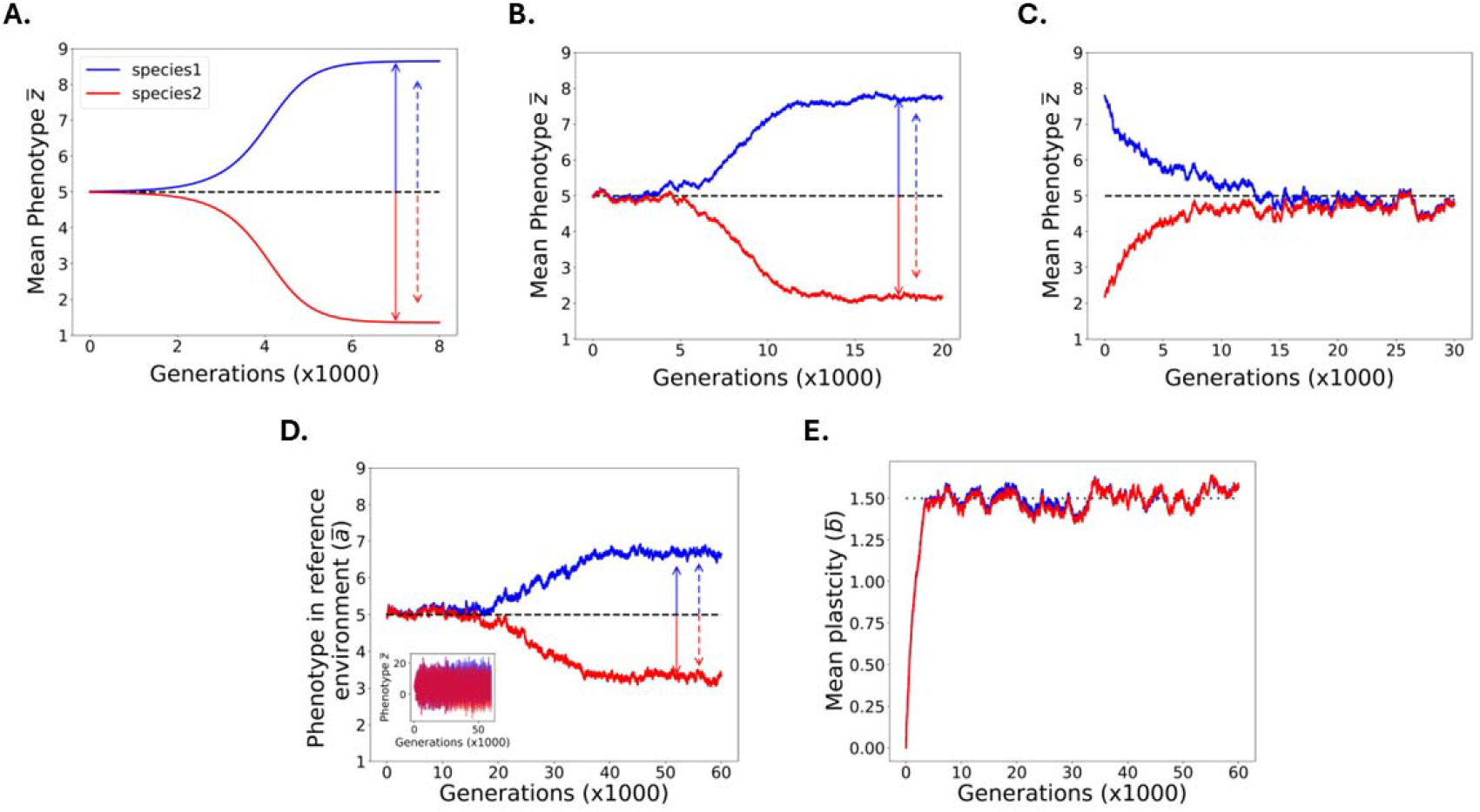
Influence of environmental fluctuations and evolution of plasticity on character displacement. The evolutionary dynamics of the mean phenotype 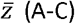 and the mean reaction norm intercept 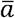 (D, with 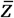 as inset) and slope 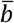 quantifying plasticity (E) are shown for individual simulations of two symmetrically competing species (represented in blue and red). The environment is constant in A but fluctuates moderately 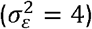in B and strongly 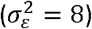 in C-E. Phenotypic plasticity is either absent and non-evolvable (A-C: *G*_*b*_ = 0), or evolvable (D-E: *G*_*b*_ =0.045). The dashed black lines in A-D represent the optimum phenotype, or its expectation in a fluctuating environment, and the dotted line in E represents the expected equilibrium plasticity, 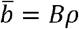. In panels A, B, and D, the vertical arrows show the expected character displacement, based either on eq. (11) combined with the mean population size computed over the last 1000 generations in the simulation (continuous arrows), or the approximation in eq. (15) that assumes the population size is at the equilibrium from eq. (13) (or eq. (17) in a fluctuating environment) (dashed arrows). All other parameter values are given in Table 1.

The criterion for CD can be expressed in a simpler form by replacing the scaled population Density *N/K* by its equilibrium value. Taking advantage of the fact that the deviation from optimum is negligible near the threshold for CD (at which 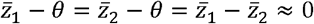), we can combine eqs (1) to (3) to get the equilibrium population size (identical in both species in the symmetric system),

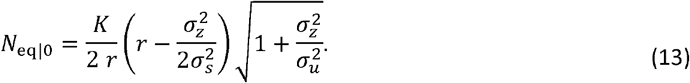

Equation (13) shows that the equilibrium population size is smaller when stabilizing selection is stronger (smaller 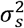), all the more as phenotypic variance 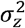 is large, because more individuals have phenotypes away from the optimum, thus reducing population growth through a standing load (Lande and Shannon 1996). In contrast, the equilibrium population size becomes larger under narrower utilization function (smaller 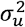).

Replacing the population density in eq. (12) by its equilibrium without CD from eq. (13) leads, after some algebra, to a much simpler criterion for CD,

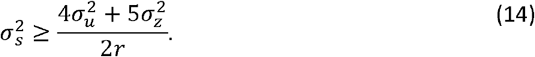

Importantly, the threshold for CD in eq. (14) only depends on the width of the utilization function, the phenotypic variance, and the intrinsic rate of increase of the species. In particular, eq. (14) predicts that the boundary of the parameter space allowing for CD is linear in the 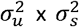 phase plane, with steeper slope when the intrinsic growth rate *r* is smaller. These predictions from are compared to intensive numerical simulations across values of 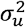 and 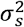 in Figure 2A.

**Figure 2:**
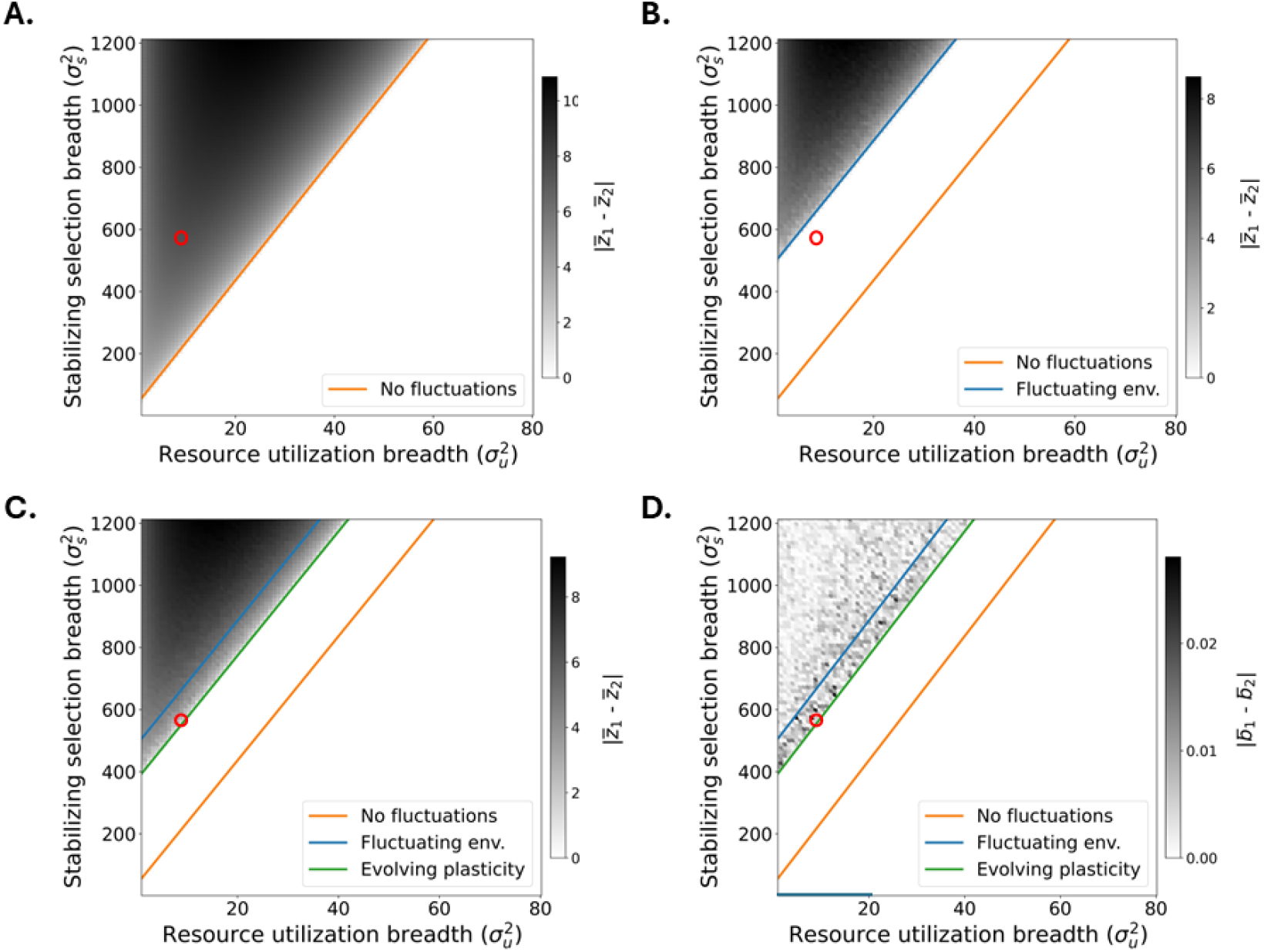
Influence of environmental fluctuations and evolution of plasticity on the conditions for character displacement. A-C. The absolute divergence in mean phenotype 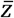 among the two competing species, averaged over 10000 generations, is shown as shading levels, for varying values of resource utilization breadth 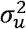 (x-axis) and stabilizing selection breadth 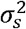 (y-axis). Simulations were performed in a constant environment with no plasticity 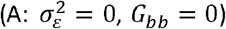 in a fluctuating environment with no plasticity 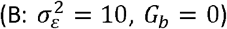 or in a fluctuating environment with evolving plasticity 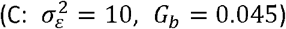. Colored lines show the analytical conditions for character displacement in these 3 contexts (eqs 14, 18, and 21 for orange, blue, and green, respectively). D: The absolute divergence in mean plasticity 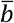 among species is shown as shading levels (note the much smaller range than in A-C) in the scenario with fluctuating environment and evolving plasticity 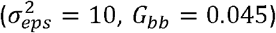 with colored lines from C reproduced for comparison. Parameter values are *A* = 5, *τ* = 0.5, and the rest is as specified in Table 1. The open red circles show the combination of 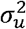 and 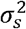 (10 and 550 respectively) used for the simulations shown in Figure 1.

Lastly, inserting the predicted population size from (13) into the character displacement in (11) yields

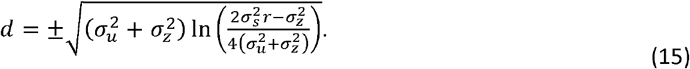

Because this prediction is based on an equilibrium population size *N*_*eq*_ that neglects CD, it is expected to underestimate the amount of CD, as relaxed competition under CD leads to higher equilibrium population size, and thus to higher CD (larger in eq. 11). This can be seen in Figure 1A, where the dashed arrows representing eq. (15) underestimate the true CD, while the prediction that inserts the actual equilibrium population size from simulations in eq. (11) (continuous arrow) matches the simulated CD.

### Environmental fluctuations can cause the collapse of CD

When a randomly fluctuating environment causes the optimum phenotype to vary over time, the evolutionary process is no longer deterministic but stocha s t ic, even in an infinite population. Below we focus on expectations under this process, denoted as ⟨ ⟩, but keeping in mind that stochasticity may lead to substantial variation around this expectation.

To understand the influence of random environmental fluctuations on the expected CD, we start by observing that fluctuations in the optimum only affect the component of fitness due to stabilizing selection 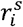 (eq. 2), not the competition term 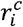 (eq. 3). In addition, the expressions for the selection gradient components in eq. (5) and (6) remain essentially unchanged by fluctuations, if we replace population sizes and mean phenotypes by their expectations. However, the expected population size (appearing in eq. (6)) is biased downwards. The reason for this bias is that environmental fluctuations add stochastic variance to the lag load caused by mismatches between the mean phenotype and the optimum. This leads to a reduction of expected Malthusian fitness (via eq. 2), which can be described as a stochastic lag load (Chevin, Cotto and Ashander 2017). Denoting as *x* the stochastic component of deviations from the optimum, with mean 0 and variance 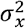, the expected stabilizing selection component of fitness is

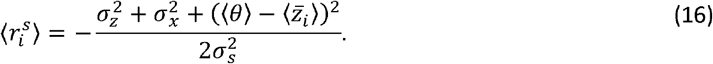

Replacing this for eq. (2) yields for the expected population size with fluctuations, in the symmetric case without CD,

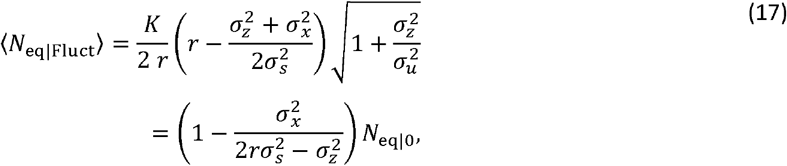

where *N*_eq|0_ is the equilibrium population size in a constant environment (eq. 13). The main influence of fluctuations in the optimum on character displacement is thus to reduce the average population size in by a proportion 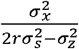 (assuming stabilizing selection is weak enough that 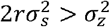). As lower population size leads to weaker competitive selection, stabilizing selection then also needs to be weaker to allow for CD. This prediction can be made more precise by inserting eq. (17) into eq. (11), leading to the criterion for CD with fluctuations

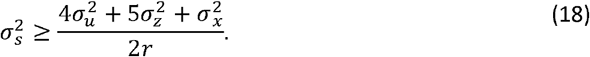

Hence, the threshold peak width 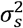 above which CD is possible increases by 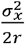 relative to the case without fluctuations, where 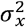 is the variance of deviations from the optimum (phenotypic mismatch). Interestingly, the criterion otherwise retains the same dependency to 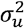, that is, threshold remains a straight line with same slope in the 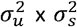 plane, but shifted upwards by 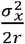. This prediction is compared to numerical simulations across values of 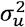 and 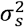 Figure 2A (blue line and symbols). In the simulations, 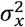 was replaced by 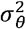, the variance of the optimum phenotype, thereby neglecting evolutionary tracking of the optimum by the mean phenotype. This will be a good approximation as long as additive genetic variance of the trait and autocorrelation of the optimum are not large; when these assumptions are violated, more complete expressions exist for adaptive tracking of an autocorrelated random optimum (Charlesworth 1993; Lande and Shannon 1996; Chevin 2013; Chevin, Cotto and Ashander 2017).

Character displacement may be preserved on average under moderate environmental fluctuations, but with reduced magnitude as illustrated in Figure 1B. However, large environmental fluctuations cause a collapse of CD, where species converge to the same mean phenotype at the average optimum ⟨ θ ⟨following the onset (or amplification) of environmental fluctuations (Figure 1C). Any combination of strengths of stabilizing and competitive selection in between the blue and orange lines in Figure 2B is susceptible to such collapse of CD under environmental fluctuations. Our results are consistent with those of Johansson (2008), who also found (albeit with simulations only, not complemented by analytical solutions) that a randomly fluctuating environment (in his case, a random walk) can lead to reduced population size and possibly extinction, altering the outcome of evolution under competitive selection.

### Evolution of plasticity can restore CD by buffering the impact of fluctuations

The influence of phenotypic plasticity can also be understood through its effect on the stochastic variance of the mismatch with optimum. With linear reaction norms and changes in the optimum with the environment (following eqs. 7-8), neglecting adaptive tracking by evolution for simplicity (but see Michel, Chevin and Knouft 2014; Tufto 2015), the variance in phenotypic deviations from the optimum is

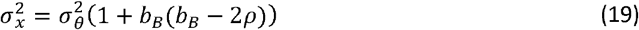

where 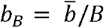 is the reaction norm slope scaled to the slope of the optimum, and *ρ* is the autocorrelation of the environment between development and selection (Chevin, Collins and Lefèvre 2013). This shows that phenotypic plasticity with the same sign as temporal autocorrelation *ρ*, and magnitude lower that |2 *ρ* |, reduces the magnitude of fluctuations in mismatch with optimum. If plasticity varies genetically, then in the long run it will evolve to match the level of environmental predictability, *b*_B_=*ρ* (Gavrilets and Scheiner 1993a; Lande 2009), such that

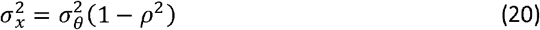

Hence, evolution of adaptive plasticity buffers the influence of a moving optimum on fluctuating selection, to an extent that depends on the predictability of selection upon development, here quantified by environmental autocorrelation *ρ*.

Combining eqs (18) and (20), the condition for character displacement with evolving plasticity becomes

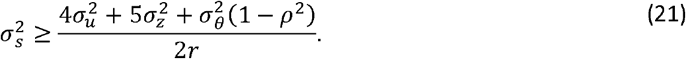

This shows that evolving plasticity can partly compensate for the detrimental effect of environmental fluctuations on CD, to an extent that depends on the predictability of selection upon development. With perfectly predictable selection (*ρ*^2^ =1) the influence of environmental fluctuations entirely vanishes because all fluctuations in the optimum can be tracked perfectly by plasticity, such that eq. (21) collapses to eq. (14).

Consider a scenario where the regime of environmental fluctuations shifts over time. While an abrupt increase in environmental fluctuations can initially lead to the collapse of CD, evolution of adaptive plasticity to match the level of environmental predictability may later restore CD, by reducing the impact of fluctuations on population dynamics and competition strength. This scenario is illustrated in Figure 1D-E, in the case where there is initially no plasticity 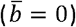, as expected in an initially fully unpredictable environment. As long as plasticity remains low (left in Fig. 1E), the mean phenotypes of both species in the mean environment (reaction norm intercept 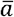) are similar, and close to the mean optimum (left in Fig. 1D). However, after plasticity has evolved in both species to its equilibrium determined by environmental predictability, 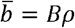 (right in Fig 1E), reaction norms Intercepts 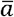, start to diverge among species, leading to phenotypic CD (right in Fig 1D). This scenari where evolution of plasticity can restore CD that was previously prevented by environmental fluctuations, occurs in the parameter range in between the blue and green lines in Figure 2C, which is larger under higher environmental predictability.

Importantly, the same level of phenotypic plasticity evolves in both species if they experience the same predictability of selection (Figure 1E). A small divergence in plasticity can be observed in the parameter space leading to phenotypic CD (Figure 2D), but with a magnitude that is negligible relative to the mean plasticity (Figure 1E). Once this equilibrium plasticity is achieved, plastic responses are essentially identical in both species. Although plasticity amplifies phenotypic fluctuations across environments (inset of Figure 1D), it does so similarly among species, such that their phenotypic divergence remains constant. This is no longer true when different levels of plasticity are favored in different species, as we now investigate.

### Competition can cause convergent character displacement in plasticity

When two competing species have slightly different mechanisms for perceiving environmental cues, or different time windows of sensitivity to these cues (because of life-history differences), then they may experience different predictabilities of selection upon induction of the plastic trait. This will cause evolutionary divergence in phenotypic plasticity in the absence of competition. However, the interplay of this process with competitive selection and character displacement is more complex.

First, when species differ in plasticity and environmental cues, the eco-evolutionary system is no longer symmetric (even when the symmetry assumptions above hold), because interspecific differences in environmental predictabilities lead to differences in expected maladaptation and population size in a fluctuating environment. Second, when species differ in plasticity (and environments of development), their mean phenotypic divergence is no longer constant, but instead changes with the environment. We thus need to distinguish character displacement in the reference environment (i.e., CD in mean reaction norm intercept 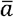) from CD in plasticity (i.e., CD in mean reaction norm slope 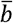). When the environment fluctuates over time, differences in plasticity cause the magnitude of competitive selection to fluctuate (because of fluctuations in phenotypic divergence, from eq. 6), which affects the expected selection gradients on the trait and its plasticity.

The conditions for CD in reaction norm intercept become more stringent when species differ in plasticity and environmental cues. As shown in the Appendix in Supplementary material, the formula for the equilibrium CD then takes a similar form as eq. (11), but with two main changes. First, the population size that matters is the average among species *N =*(*N* _1_ + *N* _2_)/2(while the symmetric model above assumed that *N* _1_ = *N* _2_ = *N*). If one of the species experiences lower environmental predictability, then its stochastic lag load will be higher (even after plasticity has evolved to its equilibrium, following eq. 20), and its expected population size will be lower. Unless this is compensated by a symmetric increase in equilibrium population size in the other species, *N*_*m*_ will be overall reduced by asymmetries introduced by differences in plasticity. If environmental fluctuations are very large, the expected population growth rate can even become negative for the species experiencing low environmental predictability, causing them to decline towards extinction (Lande and Shannon 1996; Chevin, Cotto and Ashander 2017), affecting coexistence and the scope for CD.

The second difference with eq. (11) is that 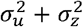 is replaced by 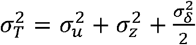,where

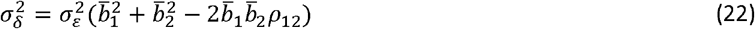

is the temporal variance in mean phenotypic divergence among species, arising from their differential plastic responses to the environmental fluctuations. The latter have variance 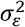,and autocorrelation *ρ*_12_ between the timings of development in species 1 and 2. As shown in eq. (22), the magnitude of fluctuations in phenotypic divergence is larger when the species differ more in plasticity, and/or have uncorrelated environments of development (small *ρ*_12_). Based on the results in the Appendix in Supplementary material, we can then define an approximate criterion for CD similar to eq. (21), leading to

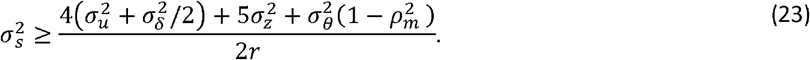

where 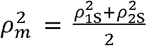 is the mean squared autocorrelation between development and selection among species. This shows that, all else being equal, stabilizing selection needs be to weaker (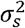 larger) for CD to exist at the phenotypic level, all the more so as environmental fluctuations are large and species differ in plasticity, such that 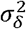 is large. When the conditions for CD in reaction norm intercept are satisfied, its relative magnitude in each species depends on the relative abundance of the other species exerting selection on it (that is, CD in species 1 is proportional to *N*_2_/ *N*_*m*_, and reciprocally; Appendix in Supplementary material).

Beyond its effect on reaction norm intercept, competition also influences the evolution of plasticity (reaction norms slope) when environmental cues differ among species. As shown in the Appendix in Supplementary material, the expected selection gradient on plasticity in species *i* is then

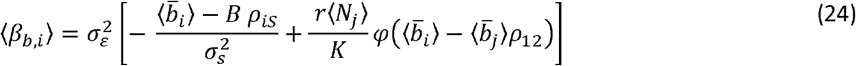

where

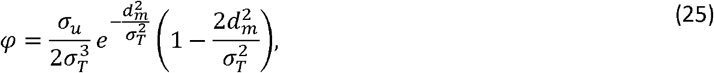

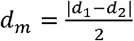 is the mean magnitude of CD (deviation from optimum) in the reference environment (equal to half the interspecific divergence in reaction norm intercept), and *ρ*_*is*_ is the correlation between the environment of development in species *i* and the environment of selection. Equation (24) first shows that selection on plasticity is overall stronger, and thus plasticity will evolve faster, when the variance 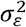 of environmental fluctuations is larger, consistent with previous theory (Gavrilets and Scheiner 1993a; Lande 2009). The terms in the brackets of eq. (24) further show how the two components of selection influence evolution of plasticity. Stabilizing selection (first term in brackets) pulls the mean plasticity in species *i* towards the slope of the optimum *B* multiplied by the predictability of selection in that species (quantified by *ρ*_*is*_), consistent with previous theory without competition (Gavrilets and Scheiner 1993a; Lande 2009; Tufto 2015). On the other hand, competitive selection (second term in brackets) causes plasticity in species *i* to evolve in response to the plasticity in the other species 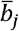 weighted by the correlation between their environments of development *ρ*_*is*._

The direction of this effect of competitive selection depends on the sign of *φ*, which itself depends on divergence in reaction norm intercept, quantified by *d*_*m*_ (from eq. 25). When *d*_*m*_ is small 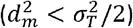, then *φ* > 0 and plasticity in species *i* is pushed a way from plasticity in the other species *j*, causing divergent CD in plasticity. However in the opposite regime where *d*_*m*_ is large 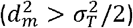 them *φ* > 0 and plasticity in species i is *pulled towards* the plasticity in species *j*. In other words, for large enough phenotypic CD in reaction norm intercept, competitive selection causes convergent rather than *divergent* CD for plasticity, whereby the reaction norm slopes of two species are more *similar* (rather than more *different*) when they compete in sympatry than if they were alone. This occurs because when species differ in plasticity and the environment fluctuates, their phenotypic divergence also fluctuates (with variance 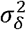, eq. 22), and this decreases the expected mean fitness (thus favoring more convergent plasticity) when CD in intercept is large, but increases it (favoring more divergent plasticity) in the opposite case. Replacing *d*_*m*_ by its expectation, the condition for competitive selection to cause convergent CD in plasticity becomes (Appendix in Supplementary material)

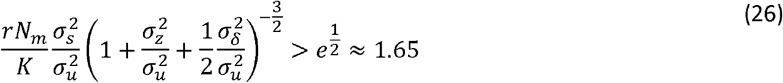

This predicts that, all else being equal, convergent CD in plasticity is more likely when (i) environmental fluctuations are small (because this leads to smaller 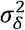, from eq. 22); (ii) stabilizing selection is weak (large 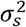) and competitive selection is strong (small 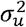); and (iii) the mean population size in both species is large (large *N*_*m*_). Conditions (ii) and (iii) coincide with those for large CD at the phenotypic level (as shown in previous sections), consistent with the fact that large (divergent) phenotypic CD in the reference environment (CD in intercept) favors convergent CD for plasticity. The net CD in plasticity under the joint influences of stabilizing and competitive selection depends on the relative magnitudes of the two components of selection, which from eq. (24) is determined by the product 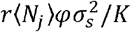; competitive selection dominates when this term is larger than 1, while stabilizing selection dominating otherwise.

These predictions are compared to simulations results in Figure 3. We considered two species that differ in their time windows of sensitivity to a single fluctuating environmental variable, thus effectively responding to distinct (but correlated) cues. As predicted by the analysis, the magnitude of environmental fluctuations has critical qualitative and quantitative impacts on CD in the trait and its plasticity. When environmental fluctuations are moderate (left panels in Fig. 3), phenotypic divergence in reaction norm intercept depends little on the between-species difference in the timing of development relative to selection (*τ*_1_vs *τ*_2_). In Figure 3A, the absolute difference in mean reaction intercept remains close to 9 regardless of *τ*_1_ and *τ*_2_ matching that without environmental fluctuations under the same strengths of stabilizing and competitive selection (compare with Fig 1A). On the other hand, the divergence in plasticity among species is strongly dependent on the difference in developmental timings. The dashed line in Figure 3B represents the predicted difference in plasticity that would evolve without competition, resulting from the difference in cue predictability (term 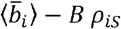 in eq. (24)). The true divergence in mean plasticity (solid lines in Figure 3B) is always below this baseline, because competitive selection leads to convergent CD in plasticity in these conditions.

**Figure 3:**
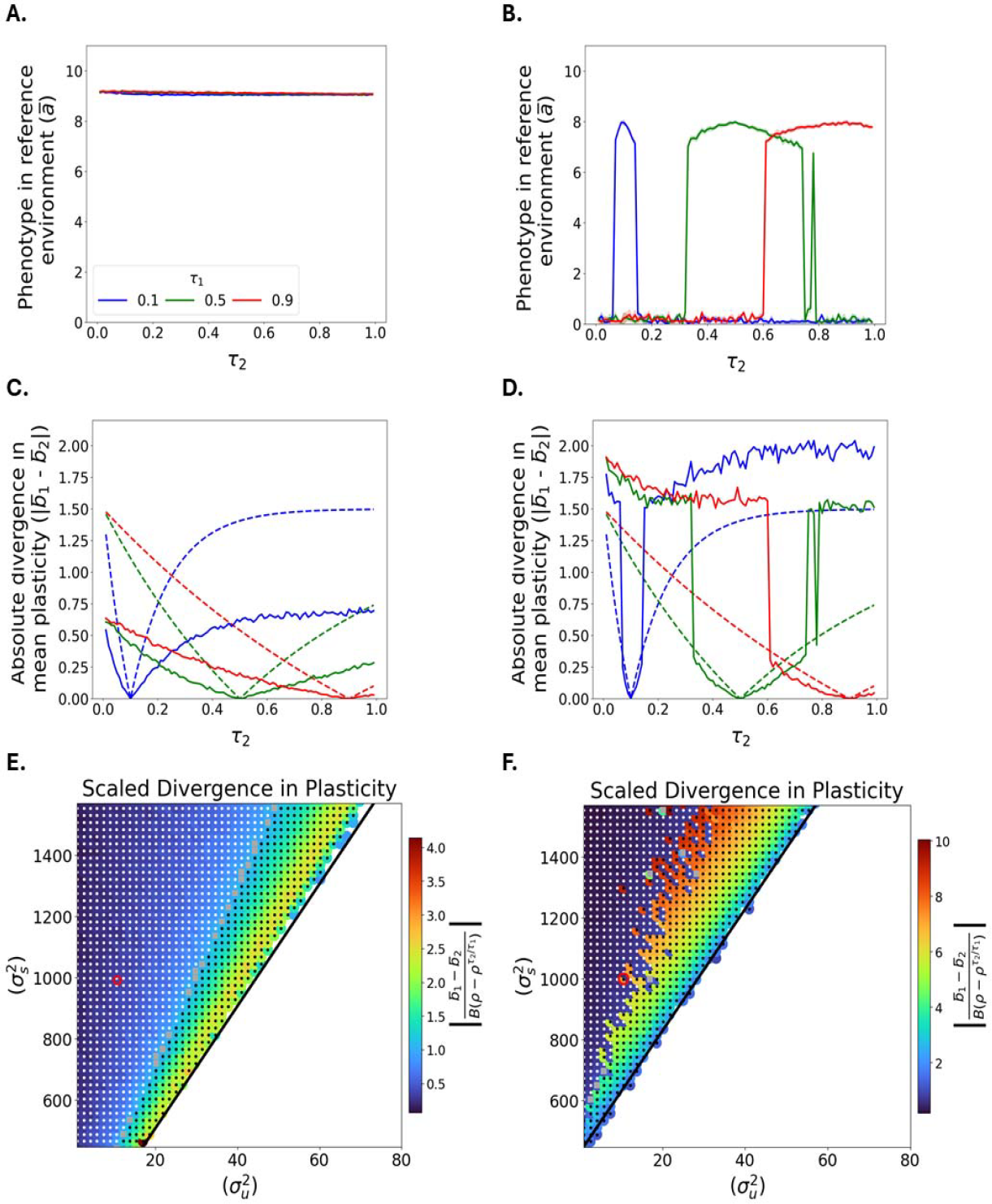
Character displacement in trait and plasticity when environmental predictability differs among species. The magnitude of between-species differences in mean reaction norm intercept 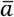 (A-B), and slope 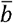 quantifying plasticity (C-D), are shown against the lag *τ*_2_ between development and selection in species 2, for several values of the corresponding lag *τ*_1_ in species 1 (colors). The dashed lines in C-D show the predicted difference in plasticity in the absence of competitive selection, which is proportional to the absolute difference in environmental predictabilities, 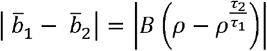 (where *ρ*=0.5) is the autocorrelation over time *τ*_1_ by definition). In the lower row (E-F), the divergence in plasticity, scaled to its expectation under no competitive selection, is shown as color levels, for varying values of resource utilization breadth 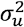 (x-axis) and stabilizing selection breadth 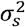 (y-axis). The developmental lag is *τ*_1_ =0.5 in species 1 and *τ*_2_ =0.4 in species 2. The predicted criterion for CD to exist at the phenotypic level is also shown as continuous black line, based on eq. (23) where 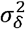 was obtained by replacing, in eq. (22), the plasticity in each species by its expectation without competition, 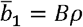 and 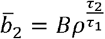 and the correlation between the environments of development in species 1 and 2 is 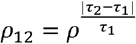. Parameter values where CD in plasticity is convergent (lower divergence than expected without competition) are marked by a white dot, those where CD in reaction norm intercept is lower than the threshold in eq. (25) (i.e., 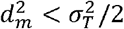 with a black dot, and those satisfying both criteria with a grey square. The variance of environmental fluctuations is 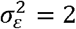 on the left (panels A, C, E) and 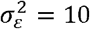 on the right (panels B, D, F). The open red circles in panels E-F show the combination of 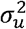 and 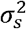 used for the simulations shown in panels A-D.

The evolutionary outcome is qualitatively changed when the variance of environmental fluctuations is much larger (right panels in Figure 3). In this context, the situation observed under moderate fluctuations (with substantial divergent CD in intercept and convergent CD for plasticity) only exists when species have similar developmental timings *τ*_*i*_ (i.e. near *τ*_2_ = 0.1 for the blue line, *τ*_2_ = 0.5 for the green line, and *τ*_2_ = 0.9 for the red line). Indeed, similar *τ*_*i*_ among species leads to a similar predictability of selection, selecting for similar plasticity, and thus moderate value of 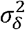 (from eq. 22), which favors convergent plasticity (from eq. 26). Conversely for more different developmental timings v (and thus larger 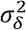), CD in reaction norm intercept collapses (Fig 3B), and divergent rather than convergent CD is observed for plasticity (solid line above dashed line in Fig 3D).

The lower panels in Figure 3 (E-F) shows how divergence in plasticity, scaled to its expectation without competition, depends on the strengths of stabilizing and competitive selection. As expected from eq. (26), weak stabilizing selection and strong competitive selection (large 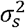 and small 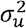, upper left corner in Figure 3E,F) lead to convergent CD in plasticity (dark blue colors in Fig. 3E-F). In contrast, smaller 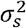 and larger 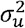 are characterized by divergent CD in plasticity (warmer colours in Figure 3 E-F). These effects can be understood through the influence of CD in reaction norm intercept, predicted by eqs (24-25). The white dots in Figure 3E-F show conditions where CD in plasticity is convergent, the black dots are where CD in intercept is lower than the threshold in eqs (24-25) 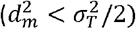, and grey squares jointly meet both these criteria. Black and white dots almost perfectly partition all the parameter space with CD (except for a small overlap with gray squares at their boundary), confirming that the magnitude of phenotypic CD in the average environment is a very good predictor of whether CD in plasticity is convergent (under large phenotypic CD) or divergent (under small phenotypic CD). The parameter regions leading to divergent CD in plasticity (warm colors in fig. 3 E-F) lies next to the boundary for CD to exist at the phenotypic level, shown as black line (based on eq. 23) in figure 3EF.

As the magnitude of environmental fluctuations increases, the parameter region leading to divergent CD in plasticity expands (Figure 3F vs 3E), but convergent CD also becomes more pronounced (darker blue in upper left corner in Figure 3F). Furthermore, large environmental fluctuations are characterized by high instability, where the system can change from divergent to convergent CD in plasticity (and concomitantly from large to no CD in reaction norm intercept) for a small parameter change, probably because of an influence of initial conditions (Fig. 3F). This instability can also by visualized in Fig. 3D, where the solid green line “flickers” as *τ*_2_ reaches large values.

The lower right corner in Figure 3E-F, with small 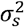 and larger 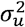, leads to extinction of at least one of the species, thus preventing computation of their divergence. This occurs because of the stochastic load imposed by a randomly fluctuating environment. Note that the parameter region leading to extinction moves up under larger environmental fluctuations (Fig. 3F vs 3E). This means that larger environmental fluctuations cause extinction even when the fitness peak is broader, as expected since the stochastic load depends on the ratio of the magnitude of environmental fluctuations 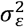 to the strength of stabilizing selection 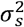 (Lande and Shannon 1996; Chevin, Cotto and Ashander 2017).

## Discussion

Both the biotic and abiotic environment contribute to the niche of a species, but how much each of them drives adaptation, and how important their interaction is for evolution, remain little understood. When the physical environment changes (e.g. rising temperature), is directional selection mostly caused by this change *per se*, or by interacting species that respond to it? And how often does adaptation to interacting species cause evolution of the fundamental niche? The intensity of ecological interactions often varies along gradients of e.g. temperature or salinity (von Humboldt and Bonpland 1807; Williams 1998; Maestre *et al*. 2009), and so will vary the selective pressures, from being driven mostly by the physical environment, to resulting mostly from interacting species. If these components of selection act on the same or on correlated traits (as implied by the notion of ecological trade-offs), then selection caused by ecological interactions will influence adaptation to the abiotic environment, even when the latter is assessed without interactors. Therefore, studies investigating environmental tolerance curves in the laboratory, without interacting species, are likely to reveal variation in fitness that partly reflects a history of adaptation to interacting species along the same environmental gradient in nature.

Ecological character displacement (CD) mediated by competition illustrates this phenomenon well. In environments where CD is most pronounced, mean phenotypes deviate from their optimum determined by the intrinsic requirements set by the environment, and thus appear as maladapted in the absence of competitors. As a result, the fundamental niche is shifted towards environments where competition has been less intense over recent evolutionary history (Slatkin 1980; Case and Taper 2000). Figure 4A illustrates this niche shift for species that evolved in a constant environment and without plasticity (as in Figure 1A). Nevertheless, we have showed that this process can be altered when the environment fluctuates over time. A randomly fluctuating optimum causes variance in phenotypic mismatch that reduces the expected mean fitness, such that the equilibrium population size is lower on average than without fluctuations (Lynch and Lande 1993; Lande and Shannon 1996; Chevin, Cotto and Ashander 2017). This leads to decreased competition intensity, tilting the balance from competitive towards stabilizing selection, and reducing the likelihood and magnitude of CD (as also shown by Johansson 2008 under a random walk, rather than the stationary fluctuations as assumed here). Figure 4B illustrates how such a collapse of CD (as shown in Fig. 1C) causes the fundamental niches of the competitors to converge. This influence of a fluctuating environment on CD, mediated by the expected population size, is similar to how environmental stochasticity can cause shifts from *K*- to *r*-selected phenotypes under density-dependent selection (Lande, Engen and Sæther 2009; Engen, Lande and Sæther 2013).

**Figure 4:**
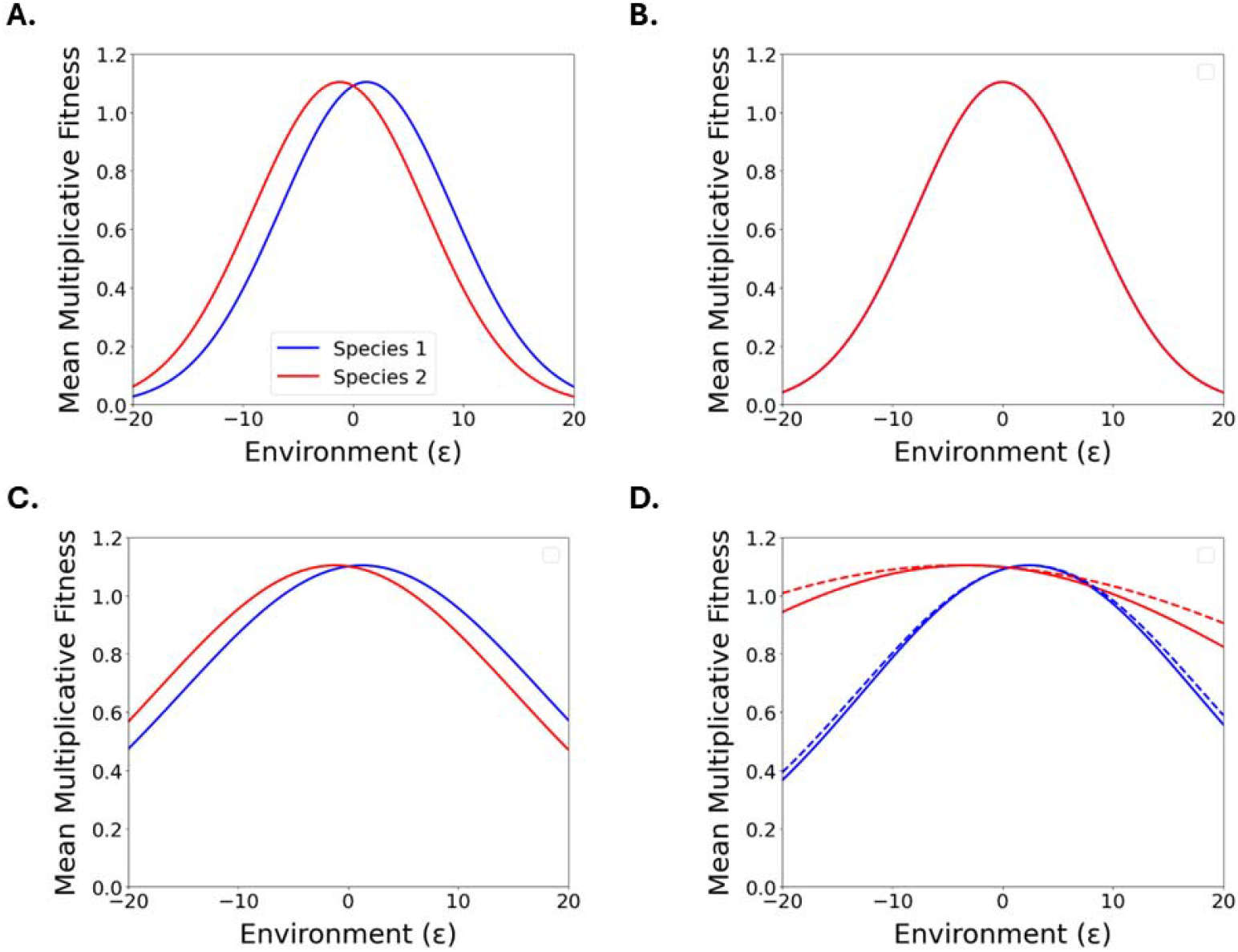
Influence of competition, environmental fluctuations, and plasticity on evolution of the fundamental niche. Environmental tolerance curves represent fitness against an environmental variable, in the absence of competitors. The mean environmental tolerance curve in a species is a representation of its fundamental niche. The mean multiplicative fitness of each species in the absence of competition, 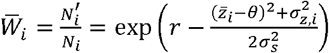 (from eq. (1)), is here plotted against the environment, for several of the scenarios investigated above. A: Conditions that lead to the evolution of character displacement in a constant environment and without plasticity (corresponding to Figure 1A) cause the niche of the two species to shift apart, such that their mean fitness is maximized in environments that differ from where they have experienced selection (ε in our simulations). B: Adding environmental fluctuations (still without plasticity) leads to the collapse of CD (same parameters as Figure 1C), such that both species converge to the same fundamental niche (species 1 in blue is not visible). C: Evolution of plasticity in a partly predictable environment (conditions similar to Figure 1D-E) broadens the fundamental niche, and here restores the niche shift caused by phenotypic CD. D: With different environmental predictabilities among species (condition corresponding to Figure 3 A,C, with *τ*_1_ = 0.9 and *τ*_2_ =0.3), divergence in phenotypic plasticity causes the niche breadths of two species to differ, in addition to their difference in niche position due to character displacement in reaction norm intercept. The dashed lines show the corresponding tolerance curves without competition, which would be both broader and more different among species.

Evolution of adaptive phenotypic plasticity can partly reverse this influence of environmental fluctuations on CD, to an extent that depends on the predictability of environmental fluctuations. As shown in Figure 4C, evolution of increased phenotypic plasticity leads to broader tolerance curves and fundamental niches, together with shifting their positions if it restores phenotypic CD. That phenotypic plasticity, when it matches the level of environmental predictability, can buffer the influence of environmental fluctuations on maladaptation and demography, has long been known theoretically (Reed *et al*. 2010; Chevin, Collins and Lefèvre 2013). A meta-analysis of breeding time in birds and mammals has confirmed that plastic responses track movements of the optimum phenotype, thus partly buffering the impact of environmental fluctuations on phenotypic mismatch (de Villemereuil *et al*. 2020). On the other hand, detailed analysis of one population of great tits in this database showed that greater phenotypic mismatch does not necessarily lead to reduced population growth, because fewer surviving offspring at this stage of the life cycle are compensated by reduced competition at a later stage, with no net demographic cost (Reed *et al*. 2013). Beyond this study case, the extent to which phenotypic mismatch (influenced by phenotypic plasticity) drives population dynamics, the strength of competition, and evolution of CD, will depend on the degree of excess fertility in a population, that is, on how much the total number offspring exceeds the requirement for replacement (as discussed in Chevin and Bridle 2025).

These demographic considerations are all the more relevant as eco-evolutionary dynamics are crucial to the processes we have investigated. Even in a constant environment, the intensity of competitive selection depends on the population densities of competing species (from eq. 6). The demographic potential of species, as measured by their intrinsic rate of increase (here expressed per generation), is thus an integral part of the condition for CD at equilibrium (eq. 14). In our simulations, we used a moderate demographic potential of *r* = 0.1,, such that both species can grow at most by ∼10.6% per generation (but usually much less because of maladaptation and/or density dependence). As a consequence, CD is only possible under very weak stabilizing selection (Figures 2-3), below what is usually reported in the literature (Kingsolver *et al*. 2001; de Villemereuil *et al*. 2020, approximating the absolute value of standardized quadratic selection gradients as 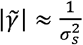), but consistent with previous theoretical findings by Goldberg and Lande (2006). However higher demographic potential (larger *r*) would allow CD to occur in the face of stronger stabilizing selection (smaller 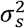). For instance with *r* = In (2) *≈* 0.69, corresponding to 1 doubling per generation in exponential phase, similar to what is observed for humans and many microbes, the threshold for CD becomes 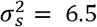 for 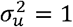 (strong competitive selection) and 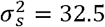 for 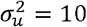 (moderate competitive selection). This leads to 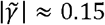 and 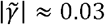 espectively, which is more in line with empirical measurements of stabilizing selection (Kingsolver *et al*. 2001; de Villemereuil *et al*. 2020).

If species experience environmental fluctuations with different degrees of predictability, for instance because their life histories differ to some extent, then they are expected to evolve different levels of phenotypic plasticity. When this occurs, their fundamental niches will differ in breadth in addition to position (if CD in reaction norm intercept exists), as illustrated in Figure 4D (for conditions corresponding to Fig. 3A,C). However, an unexpected outcome of our analysis was that competition that favors more different phenotypes between species, as commonly assumed in models of character displacement (Slatkin 1980; Taper and Case 1992; Doebeli 1996b; Goldberg and Lande 2006), can lead to *convergent* rather than *divergent* phenotypic plasticity, where reaction norm slopes are more *similar* among competing species than if they were alone (continuous vs dashed lines in Fig. 3C, and in Fig. 4D for tolerance curves). That character displacement can lead to phenotypic convergence has been highlighted previously (Grant 1972; Abrams 1987; Fox and Vasseur 2008), but not with respect to phenotypic plasticity under a simple scenario of competition for nutritionally substitutable resources. While this result seems paradoxical, it is in fact meaningful in hindsight. When species differ in phenotypic plasticity, their phenotypic divergence varies across environments. Such plastic variation in CD was not specifically favored in the simple ecological scenarios we explored (neither was it in previous work on ecological CD). With environmental fluctuations, differences in plasticity cause an additional term in expected mean fitness and selection gradients, which favors more similar plasticity among species when their phenotypic CD in the average environment is large (eqs. 24-25). This effect may however be reversed in more complex ecological scenarios that might favor variation in phenotypic divergence among environments. This could occur when the balance between competitive and stabilizing selection varies across environments, for instance because of a change in carrying capacity (see eqs 4-6 and 11). We have briefly explored this type of scenario (not shown), but have not been able to find conditions cancelling convergent CD in plasticity, so we leave a more thorough investigation of this question to future studies.

In line with earlier theory on CD (Case and Taper 2000; Goldberg and Lande 2006; Goldberg and Price 2022), we have treated additive genetic variance as an input parameter rather than as an output of the model, which has a number of implications. First, we have assumed constant additive genetic variance within a given simulation (and in analytical results), but genetic variance would change over time with more explicit genetics. In particular, genetic variance is likely to increase when selection changes from stabilizing to disruptive, as species approach the same optimum and start competing more intensely. This process is unlikely to qualitatively alter our conclusions, but would we worthwhile investigating with individual-based simulations in future studies. Second, we have assumed for simplicity that the genetic variances are identical in both species (as done by Case and Taper 2000; Goldberg and Lande 2006; Goldberg and Price 2022). This leads to conditions for CD that do not depend of the amount genetic variance (eqs. 14, 18, 21, 23), but in a more realistic setting, more complex expressions could be obtained that would depend of ratios of genetic variances. Third, the equilibrium genetic variance could vary across simulations that differ in the strengths of stabilizing and competitive selection (in Figures 2-3) if the mutation parameters (rate and phenotypic effects) were held constant, as commonly done in individual-based simulations. However the additive genetic variance is more directly amenable to empirical measurement than mutational parameters, making it meaningful to use the former as basic parameter the model, as we did. Fourth, with genetic variance in plasticity, environmental fluctuations lead to fluctuations in additive genetic variance and heritability, bearing on responses to selection and eco-evolutionary dynamics (Lande 2009; Tufto 2015; Ashander, Chevin and Baskett 2016). The latter effect was studied in detail (without competition) in earlier theory (Lande 2009; Tufto 2015; Ashander, Chevin and Baskett 2016), so we did not explore it further here, but it was included in our simulations.

Other extensions would be worthwhile investigating. In particular, we assumed that (i) a single trait mediates both competition and adaptation to the abiotic environment, in line with earlier models of CD (Slatkin 1980; Case and Taper 2000; Goldberg and Lande 2006; Goldberg and Price 2022); and (ii) a single environmental variable influences both the position of the optimum and plasticity, in line with most models of plasticity (de Jong 1990; Gavrilets and Scheiner 1993a; Lande 2009; Goldberg and Price 2022). Regarding the latter, temperature for instance is known to induce plasticity in a number of traits, and to also affect phenotypic selection, sometimes on the same traits. For instance breeding time in hole-nesting birds responds plastically to temperature, and has an optimum that also depends on temperature (Charmantier *et al*. 2008; Reed *et al*. 2013; Chevin, Visser and Tufto 2015). The timing of breeding should also impact the intensity of resource competition, since an enormous amount of caterpillars are required to feed each chick until fledgling (Visser, Holleman and Gienapp 2006), such that synchronicity with other individuals or species should increase competition strength. Beyond this example, it is unclear to what extent the same traits are likely to mediate competition and adaptation to the changing environment in general. Extending the model to include two genetically correlated traits, one of which mediates competition while the other is under stabilizing selection, would be straightforward in principle, but this would require finding a multivariate definition of character displacement, or using other criteria based on e.g. invasion analysis (Germain *et al*. 2018).

Another interesting extension would be to allow for plasticity in response to the type of consumed resources, or to the presence/abundance of competitors, as shown experimentally by Pfennig and Murphy (2000). If plasticity in response to the abiotic environment and to competitor presence both vary genetically, then multivariate cues for plasticity can evolve (Chevin and Lande 2015). For instance, the abiotic environment could become an indirect cue for the presence of competitors, or conversely plasticity could evolve to be mostly in response to competitor presence, without an influence of the abiotic environment. Exploring how these scenarios influence the onset and magnitude of character displacement would further improve our understanding of the role of competition on evolution of the fundamental niche.

## Acknowledgements

We thank Ophélie Ronce and two anonymous reviewers for useful comments on this manuscript.

## Appendix: Details of derivations

### Equilibrium character displacement in a constant environment

From the evolutionary recursion in eq. (4), in the symmetric case (where indexes for species can be dropped for simplicity), denoting as *d* the deviation from optimum in species 1 (and thus −*d* in species 2), the evolutionary equilibrium satisfies

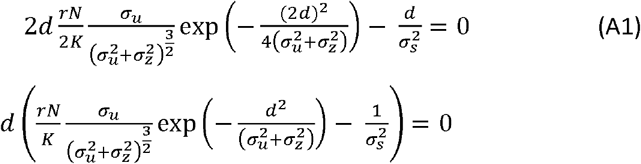

An equilibrium with *d* ≠0 exists when

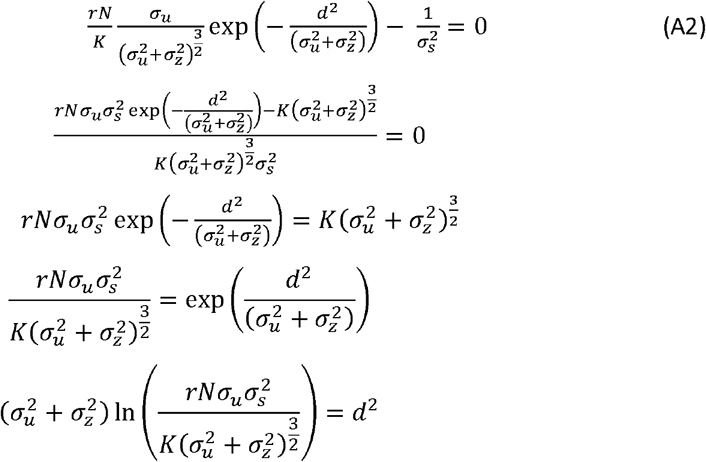

The criterion for the logarithm to be positive is

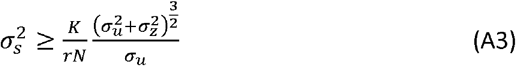

Near the threshold for CD, in the symmetric system the equilibrium population size satisfies

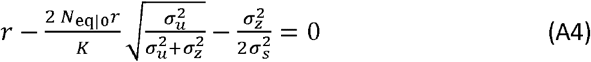

Solving yields the equilibrium population size in eq. (13) in the main text, which can then be replaced in the criterion for CD (eq. 12) to yield

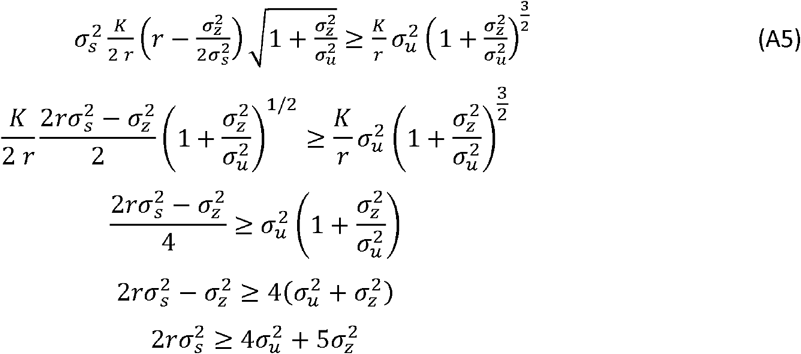

### Equilibrium character displacement in a fluctuating environment

In the presence of fluctuations, near the threshold for CD in the symmetric system the expected equilibrium population size satisfies

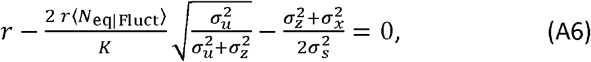

where ⟨ ⟩ denotes an expectation over the stochastic process of environmental fluctuations.

Rearranging yields

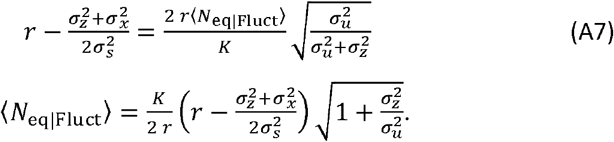

The ratio of the equilibrium population size with to without fluctuations is

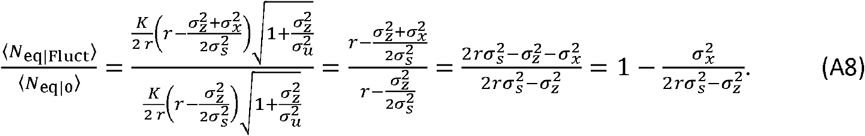

Note that the terms containing 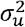 cancel out, so the expected proportional reduction in equilibrium population size caused by fluctuations is the same, whether or not density regulation is mediated by phenotype (ie, with or without competitive selection).

When CD exists under a fluctuating optimum, its expected value is

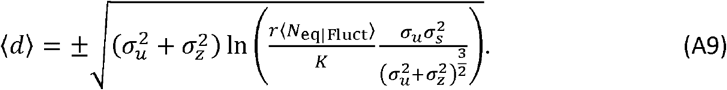

Replacing the equilibrium population size by its expectation under no CD, the term in the logarithm can be simplified as

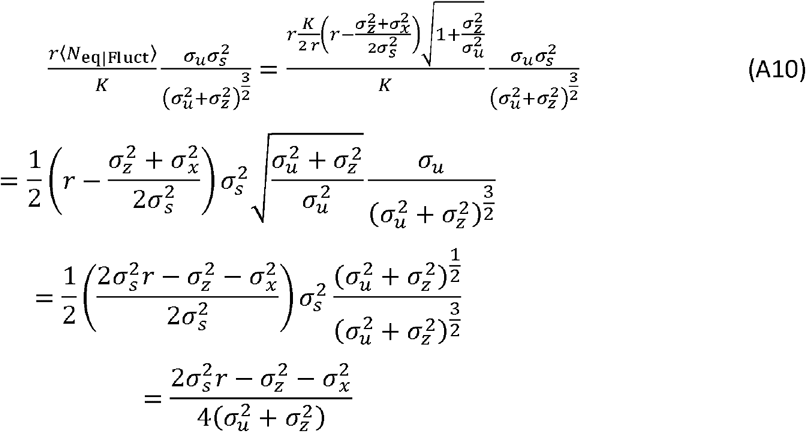

This leads to the approximate formula for the expected equilibrium CD with environmental fluctuations,

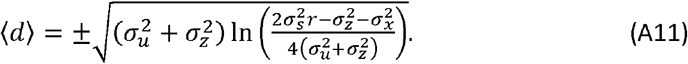

This formula is only an approximation because it is based on an expression for the equilibrium population size that assumes no CD. However, it can be used to derive the criterion for non-zero (expected) CD with fluctuations, which needs to satisfy

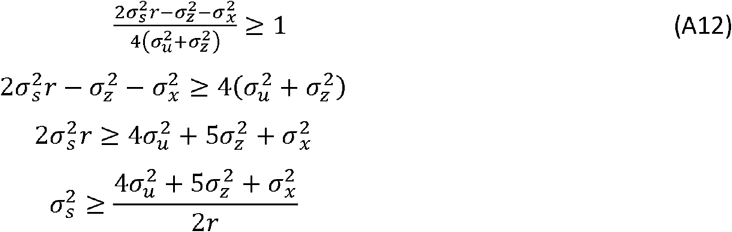

With plasticity, and neglecting adaptive tracking of the optimum by evolution of the mean phenotype, the variance in phenotypic mismatch is

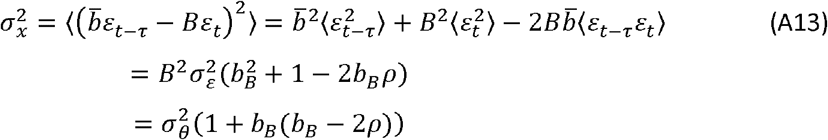

where 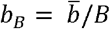 is the reac tion norm slope (plasticity) scaled to the slope of the optimum. Without plasticity, this collapses to 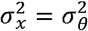.

### Divergence in plasticity

When the competing species are allowed to respond to the environment at different times (because of e.g. differences in life history), and may thus evolve different reaction norms slopes, the conditions for character displacement at the phenotypic level are modified. We first note that the symmetry assumption used above breaks down in this context, because species experiencing different environmental predictabilities and with different plasticities will also have different equilibrium population sizes and phenotypic deviations from the optimum.

The total selection gradient on expressed trait *z* in species *i* in a given generation becomes (from eqs 4-6)

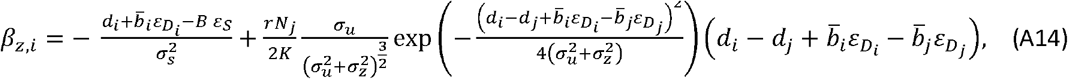

Where 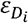 is the environment influencing plasticity for species (and similarly for 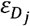 and species *j*), and *d*_*i*_ is the character displacement of reaction norm intercept in species *i*, that is, its phenotypic deviation from the optimum in the reference environment (where ε =0 by convention). The character displacements of reaction norm intercepts *d*_*i*_ and *d*_*j*_ may have different magnitudes due to the asymmetry induced by the different environmental predictabilities, but they have opposite signs, so we assume (without loss of generality) that *d*_1_ is positive and *d*_2_ negative, such that the magnitude of divergence between species in the reference environment is *d*_1_ −*d*_2_. We define the half divergence in reaction norm intercept (or mean CD in reference environment) as 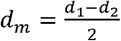. Assuming that the environments of selection ε_*S*_ and 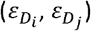 are multivariate normally distributed, and integrating the gradient over this distribution leads to the expected gradient on the trait

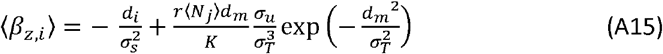

where ⟨ ⟩ denotes an expectation over the stochastic process of environmental fluctuations as before, and

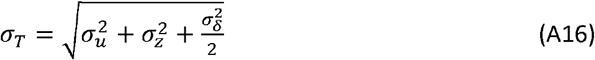

where

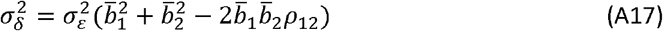

is the variance of fluctuations in phenotypic divergence among species, and *ρ*_12_ is the correlation in environments of development between species. The case where both species have the same plasticity 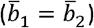 and respond to the same environment (*ρ*_12_ = 1) is recovered for 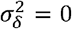, as expected. Equations (A15-A17) shows that the main effect of divergence in plasticity and differences in environments of development between species on phenotypic selection is to add half the temporal variance of phenotypic divergence 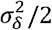 to the sum 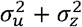.

The expected character displacements for the expressed phenotypes in both species is obtained by jointly solving ⟨ *β*_*z*,1_ ⟩ = 0 and ⟨ *β*_*z*,2_ ⟩ = 0 for *d*_1_ and *d*_2_ yielding

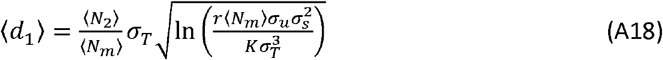

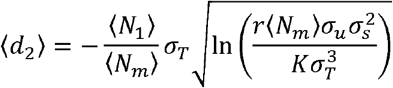

where 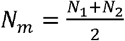 is the mean population size across species, and ⟨ ⟩ denotes a stochastic expectation as before. This shows that in such asymmetric conditions, what determines the relative magnitudes of CD in each species (when they exist) is simply their ratio of equilibrium population sizes, 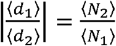 In other words, the species with the largest population size at equilibrium causes the largest CD in the other species. In addition, the magnitude of CD increases with the average ⟨ *N*_*m*_⟩ as well as with the compound competition parameter *σ*_*T*._ Provided that both species can persist (coexistence), the condition for character displacement in both species is that the term in the square root is positive, leading to

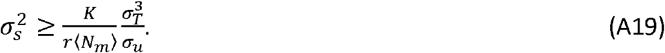

This shows that a difference in plasticity between species makes the conditions for CD at the phenotypic level more stringent, for two reasons. First, differences in plasticity 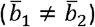 and/or in so as the variance of environmental of development (*ρ*_12_ <1) between species lead to 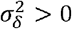 increasing *σ*_*T*._ all the more so as the variance of environmental fluctuations 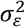 is large (from eqs. A16-A17). And second, asymmetric maladaptation can lead one population size to be well below carrying capacity, thus decreasing the mean population size across species *N*_*m*,_ if not compensated by an increase of same magnitude in the other species (which is unlikely because of density dependence). Therefore, weaker stabilizing selection (larger 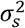) is required for CD to exist.

Divergence in plasticity is also influenced by competition in this context. To understand how, we start from the selection gradient on plasticity for species (using eq. 9)

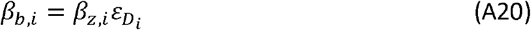

Combining with the expression for *β*_*z*,1_ above and integrating over the distribution of environments of selection and development of both species leads to the expected selection gradient on plasticity for species *i*,

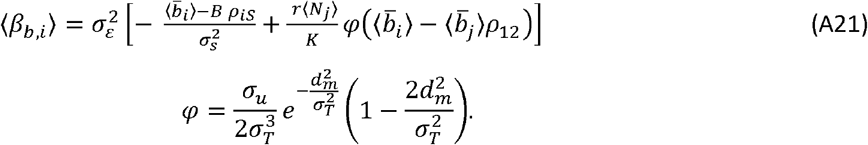

Where *ρiS* is the correlation between ion in species *i* and the environment of selection. We can make more analytical progress by replacing the CD in reaction norm intercept by its predicted expression (eq. A18), to get

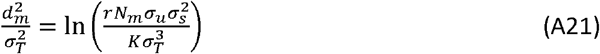

such that

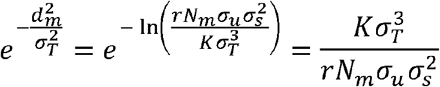

leading to

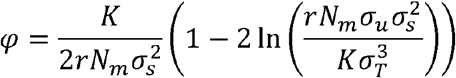

and finally

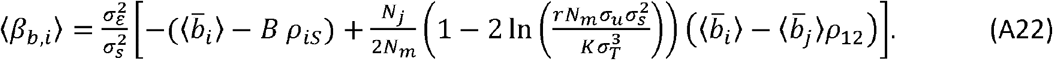

With this criterion, competition causes convergent CD in plasticity when

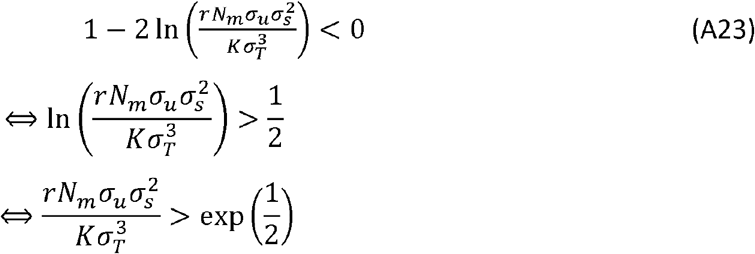

and divergent CD in plasticity otherwise. Replacing *σ*_*T*._ by its expression in eq. (A16) leads to

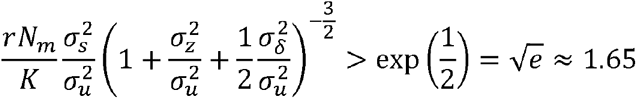

as condition for convergent CD in plasticity.

## Notes

### Competing Interest Statement

The authors have declared no competing interest.

### Summary of Updates

This version includes a new figure (Fig 4) on the evolution of fundamental niches, as well as additions to the introduction and discussion.

